# Reconstruction of the urinary tract at the appropriate time reduces fibrosis of the metanephros in rats as judged by imaging

**DOI:** 10.1101/2020.03.20.000273

**Authors:** Kotaro Nishi, Takafumi Haji, Takuya Matsumoto, Chisato Hayakawa, Kenichi Maeda, Shozo Okano, Takashi Yokoo, Satomi Iwai

**Affiliations:** Laboratory of Small Animal Surgery 2, School of Veterinary Medicine, Kitasato University, Towada, Aomori, Japan; Division of Nephrology and Hypertension, Department of Internal Medicine, The Jikei University School of Medicine, Minato-ku, Tokyo, Japan; Meiji University International Institute for Bio-Resource Research, Kawasaki, Kanagawa, Japan

## Abstract

Chronic kidney disease leads to high morbidity rates among humans. It is a serious disease that requires curative treatments other than kidney transplantation. Recently, we successfully established the iPS-derived generated kidney, which might produce urine. The urine can be directed to the native bladder with a stepwise peristaltic ureter system, followed by anastomosis with the recipient ureter for reconstruction of the urinary tract. However, the growth of the regenerated kidney varies significantly, whereas the time window of the anastomosis is quite narrow. Therefore, this study was conducted to evaluate the growth of transplanted metanephros with bladder periodically and noninvasively using computed tomography and ultrasonography. Ultrasonographic findings showed high correlations with computed tomographic findings and clearly evaluated metanephros with bladder. We found that the degree of growth of the metanephros with bladder after the transplantation differed in each individual. However, most of them reached the appropriate period for urinary tract reconstruction within 3 weeks after transplantation. Optimizing the stepwise peristaltic ureter system anastomosis by ultrasonography reduced long-term tubular dilation of the metanephros, thereby decreasing fibrosis caused by transforming growth factor-β. This may be significantly related to long-term maturation of fetal grafts. These results provide new insights into transplanting regenerated kidneys in higher animals. We are one step closer to the first human trial of kidney generation.

## Introduction

The morbidity rate of end-stage renal disease (ESRD) remains high. Although kidney transplantation is the main curative treatment, securing an available donor is difficult [1]. Apart from the number of people waiting for a kidney transplant, the number of patients undergoing hemodialysis is increasing. There is a large number of ESRD patients worldwide; furthermore, all these associated factors cause huge medical expenses and impose heavy burden on families in particular and the society in general [2].

There is a need treatment for alternatives to kidney transplantation and dialysis, which must be a fundamental treatment. There has been an attempt to make a kidney from pluripotent stem cells de novo. Takasako et al. [3,4] examined a method for producing renal organoids in vitro by aggregating nephron progenitor cells and ureteric bud. Organoids include differentiating nephrons, interstitial, and vasculature, which have matured in a culture. In addition, the use of human stem cells has made it possible to produce organoids that resemble human fetal kidneys. However, the size of organoids was smaller than 2 mm, and could not gain enough ability for the production of urine, such as functional maturation of tubules, glomerular neovascularization, and urinary excretion pathway.

To overcome these issues, we made an attempt to generate a kidney from induced pluripotent stem cells (iPS cells) using nephrogenic niche of xeno-animal as the scaffold to generate the kidney [5,6]. In this system, iPS cell-derived nephron progenitor was injected into the nephrogenic zone of xeno-embryo and cultured in the nephrogenic environment. We confirmed that injected cells continued to develop further to form a nephron, and it started producing urine following transplantation in vivo [7]. By eliminating the native nephron progenitor cells (NPCs) in the nephrogenic zone during development using genetic manipulation, pure nephron from external NPCs can be successfully generated [8]. We also confirmed that this system can generate interspecies chimeric nephron between rats and mice [9], and also iPS cells from hemodialysis patients can be used without deterioration compared with those from healthy controls (Tajiri Sci Rep). Based on this success, we are currently conducting the scale up experiment using bigger animals to proof the efficacy and safety for human clinical use. However, one big hurdle remains for the next stage. Transplantation alone does not provide a route for excretion of the produced urine. Thus, metanephros can cause hydronephrosis and renal insufficiency [11,12]. This may be solved using Stepwise Peristaltic Ureter (SWPU), which (S1 Fig.) comprises anastomosis of the ureters of the recipient rats to the bladder using a developed metanephros with bladder (MNB) [11]. This new method made it possible to continuously excrete urine produced from the MNB to the recipient bladder via the recipient ureter [11]. However, the timing of anastomosis with the ureter of the recipient after transplantation is crucial and owing to individual differences in the growth of MNB, hydronephrosis may occur at ambiguous anastomosis times [11,13]. Postrenal nephropathy due to hydronephrosis imposes a heavy burden on the kidneys, and the delayed release of obstruction has substantial effects on the kidneys [14–16]. Ureteral primordia obstruction during the fetal stage has been shown to cause dysplastic metanephros [17]. Therefore, we believe that early released obstruction is significantly involved in subsequent renal functions, even with fetal-derived grafts. In the case of xenotransplantation and MNB transplantation in large experimental animals, the effects of individual differences are considered to be greater. Appropriate time must be allowed for urinary tract reconstruction using a minimally invasive method that can be used to observe MNB over time and can be clinically performed. Clinical and general imaging methods, including contrast-enhanced computed tomography (CT) and particularly, ultrasonography have been used because they help to easily determine the condition of the body and are less invasive.

Therefore, in this study, we aimed to establish the appropriate time index for urinary tract reconstruction using morphological and histopathological examination and image analysis.

## Materials and methods

Experiment 1 was a morphological assessment of two transplanted MNBs using contrast CT and ultrasonography examinations. Experiment 2 investigated the hypothetical conditions for appropriate timing of urinary tract reconstruction based on the results of Experiment 1 when the following two conditions were met:

1. Neither hydronephrosis of the MNB metanephros, nor two or more vacuoles were observed in MNB by ultrasonography.
2. MNB bladder was larger than 0.016 cm^3^, when assessed by ultrasonography.

In Experiment 3, anastomosis was performed and the MNB grew up to 8 weeks after transplantation; then, glomerular filtration rate (GFR) measurements and histopathological examinations were performed.

### Animals

The animal rearing management was carried out according to the Kitasato University Faculty of Veterinary Medicine Animal Experiment Guideline and Manual for Rearing and Management of Experimental Animals (Approval No: 17-127, 18-127, 19-085). The rats were housed in cages under temperature and light-controlled conditions in a 12-hour cycle and were provided with fresh food and water *ad libitum*.

In Experiment 1, we used three pregnant female Lewis rats on gestation day 15 (E15) (Japan Charles River Laboratories, Kanagawa, Japan) to obtain fetal MNB. As recipient rats (organ recipient animals), we used 12 male Lewis rats (Japan Charles River Laboratories) aged 11 weeks, with a body weight of 309.0 ± 11.4 g.

In Experiment 2, we used three pregnant female Lewis rats on E15 (Japan Charles River Laboratories), and as recipient rats, 18 male Lewis rats (Japan Charles River Laboratories) aged 9 weeks, with a body weight of 243.0 ± 7.5 g.

In Experiment 3, we used three pregnant female Lewis rats on E15 (Japan Charles River Laboratories), and as recipient rats, 9 male Lewis rats (Japan Charles River Laboratories), aged 10 weeks, with a body weight of 292.8 ± 7.6 g.

### Isolation and grafting of MNB

The surgery was performed by an experienced surgeon specialized in microsurgery. Pregnant rats were anesthetized by 2.5% isoflurane inhalation. Embryos (E15) were harvested, and the pregnant rats were then killed immediately by an infusion of pentobarbital (120 mg/kg). All the embryos were euthanized by decapitation. The MNBs were dissected under a surgical microscope, as previously described [11].

### Method of MNB transplantation/urinary tract reconstruction

#### Experiment 1: Usefulness of CT examination and ultrasonography, and image evaluation of MNB

The flow of the experiment is shown in Fig. S2 A. Anesthesia was introduced and maintained in the recipient rats using 2.5% isoflurane. After performing a laparotomy through a midline abdominal incision, the intestinal tract was pulled out of the body, and the retroperitoneum and the abdominal aorta were exposed. A small incision was made to the retroperitoneum under a surgical microscope, and the first MNB (MNB1) was transplanted to the retroperitoneal space near the abdominal aorta. After transplantation, a single interrupted suture was made on the retroperitoneum using 6-0 non-absorbable suture thread (PROLENE^®^, Johnson and Johnson K.K., Tokyo, Japan). The wound was closed using conventional methods. The animals were divided into two groups. The first group comprised randomly selected rats that had the left recipient kidney removed 4 weeks after MNB1 transplantation and underwent urinary tract reconstruction by anastomosing the recipient ureter to the MNB (n = 5: anastomosis group). The second group consisted of randomly selected rats that did not undergo urinary tract reconstruction by anastomoses (n = 7: non-anastomosis group). In addition, both groups had the second MNB (MNB2) transplanted in week 4. We distinguished MNB2 from MNB1 by transplanting MNB2 to the head side of MNB1. Eight weeks after MNB1 transplantation, both MNB1 and MNB2 were removed, and histopathological examinations were conducted.

#### Experiment 2: Establishment of an appropriate anastomosis time for each MNB

The flow of the experiment is shown in Fig. S2 B. Only a single MNB transplantation was performed, with the same methods as in Experiment 1. MNB observations were performed every other day from 2.5 weeks after transplantation, and the morphological characteristics and MNB volume were assessed in the same technique as in Experiment 1. If MNBs met the conditions of appropriate timing for urinary tract reconstruction, as established in Experiment 1, the animals were euthanized and the MNB was removed. Animals that did not meet the conditions were observed until Week 5, before having the MNB removed. The removed MNB was fixed for histopathological examinations.

#### Experiment 3: MNB evaluation after SWPU at the appropriate time for anastomosis

The flow of the experiment is shown in Fig. S2 C. Only a single MNB transplantation was performed, under the same methods as in Experiment 2. MNB observations were performed every other day from 2 weeks after transplantation, and the morphological characteristics and MNB volume were assessed with the same technique as in Experiment 1. Six animals were randomly selected and if MNBs met the conditions of appropriate timing for urinary tract reconstruction, as established in Experiment 1, they underwent SWPU (n = 6: SP group). The remaining three randomly selected animals underwent SWPU at 4 weeks and were observed for 8 weeks after the transplantation (n = 3: 28UR group). After measuring the GFR, the MNBs of rats were removed and fixed for histopathological examinations. The rats were euthanized after removal of the MNBs.

### MNB assessment using contrast CT scans

For the imaging, we used the Auilion 16 Multi slice CT system (Toshiba Medical Systems K.K., Tochigi, Japan). The experiments were conducted in dynamic CT mode, at a tube voltage of 80 kV, tube current of 150 mA, imaging rotation speed of 0.5 sec/rotation, and slice thickness of 0.5 mm. The radiation exposure dose was kept unified at 392.7 mGy. The animals were anesthetized and maintained with 2.5% isoflurane at 2.5 weeks and 3.5 weeks after MNB2 transplantation (6.5 and 7.5 weeks after MNB1 transplantation, respectively), and they were all kept in the supine position during imaging.

For contrast CT examinations, the animals received a bolus injection of 0.3 mL/head Iohexol (Omnipaque^®^ 300 injection, Daiichi Sankyo K.K., Tokyo, Japan), an iodine contrast agent, into the tail vein and images were taken 30 min later, when the contrast agent was thought to have accumulated in the MNB.

After reconstitution and reconstruction of the images, we processed them using the DICOM viewer software OsiriX, to assess the morphological characteristics and MNB volume.

### MNB assessment using ultrasonography

Three technicians, who use the ultrasonography device on a daily basis, performed the examination to randomly selected rats. To carry out these tests, the rats were anesthetized and maintained until the end of the procedure using 2.5% isoflurane. Their abdomens were shaved, and the animals were kept in the supine position. The ultrasound device LOGIQ S8 (GE Healthcare Japan K.K., Tokyo, Japan) was used. After identifying the MNB through the observation of axial, sagittal, and coronal cross-sections, the maximum long-axis length of the sagittal cross-section (L), maximum short-axis width of the axial cross-section (W), and height of maximum depth (H) were measured. The volume (*V*) of the MNB bladder was also assessed.

As a probe, we used a 3-11 MHz linear array. We used the color Doppler mode to identify the presence/absence of blood flow to the transplanted MNB. To measure the MNB or MNB bladder volume, we used B mode and set the gain and depth to 90 and 2.3–2.5 cm, respectively, and then we observed the morphology and measured the size. Volume calculation by ultrasonography assumed the MNB to be a spheroid, substituting the values into the formula shown below for the volume of a spheroid:

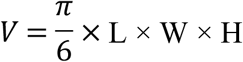

### Removal of the transplanted MNB

After anesthetizing the rats with 2.5% isoflurane, we made a midline abdominal incision and removed the MNB, which was used for histopathological examinations. After we intraperitoneally administered 125 mg/kg of pentobarbital and confirmed the cardiopulmonary arrest 15 min later.

### Histopathological examination

The MNB tissues were fixed using 4% paraformaldehyde phosphate buffer solution and were embedded in paraffin as previously described [11]. Hematoxylin-eosin (HE) dye and Masson’s trichrome staining (MT) were used in Experiments 1 and 2. In Experiment 3, the tissues were thinly sliced to 2 μm and stained for TGF-*β*1 (sc-130348: Santa Cruz, CA), collagen-α1 type 1 (sc-293182: Santa Cruz, CA), vimentin (422101: Nichirei Biosciences Inc., Tokyo, Japan) and cell apoptosis for immunostaining. TUNEL assay was performed to detect apoptotic cell death using the in situ Apoptosis Detection Kit (Takara Bio Inc., Shiga, Japan) according to the manufacturer’s instructions. The entire metanephros cut at maximum length were observed at 400× magnification, HE, MT, TGF-*β*1, collagen-α1 type 1, vimentin with 20 taken, TUNEL with ten images taken for each slide. The images were assessed by researchers, who were blinded, using the image analysis software ImageJ^®^ (National Institutes of Health, Bethesda, Maryland, USA); HE-stained slides were used to evaluate tubular lumen area and MT dyed slides to evaluate interstitial fibrosis in the metanephros.

### GFR measurement

GFR was measured using a commercially available kit (Diacolor_®_ Inulin, TOYOBO CO., LTD., Osaka, Japan). The measurement was performed according to the kit method. Eight weeks after transplantation in Experiment 3, bilateral nephrectomy was performed under the same anesthesia as mentioned previously. The blood was collected from the tail vein of the rats at one and two hours after inulin administration, and the measurement was carried out using the plasma. Normal kidney GFR values were determined by measuring healthy adult rats (Table S).

### Statistical Analyses

The results are presented as mean ± standard deviation. All statistical analyses were performed with EZR (Saitama Medical Center, Jichi Medical University, Saitama, Japan) [18]. More precisely, it is a modified version of R commander, designed to add statistical functions frequently used in biostatistics. Scatter plot and Pearson’s product ratio correlation coefficient were applied for volume comparison and storage volume comparison using ultrasonography and contrast CT examination, and the relationship between TGF-β1 expression levels and the percentage of apoptotic cells. We used the paired student’s t-test to examine the differences in volume observed over time by ultrasonography. The Mann–Whitney U test was used to analyze tubular dilation, and metanephros fibrosis was examined through histopathological examinations. A *p*-value of 0.05 was set as statistically significant.

## Results

### Experiment 1

#### MNB detection rate by contrast CT and ultrasonography examinations

Contrast CT and ultrasonography examinations allowed us to evaluate all MNBs by Week 3 (Fig. 1). However, it was not possible to assess the MNB bladder and MNB that underwent hydronephrosis. Ultrasonographic examinations allowed us to detect MNB that was present near the aorta from the early stage, and the margins were clear too. Furthermore, it allowed 100% recognition of both MNB1 and MNB2, from Week 3 onward after transplantation (Fig. 2) and partial observation of the newly formed blood vessels around the MNB by the color Doppler method (Fig. 1 D). Additionally, it was possible to confirm the metanephros (Figs 1B and C).

**Fig. 1.**
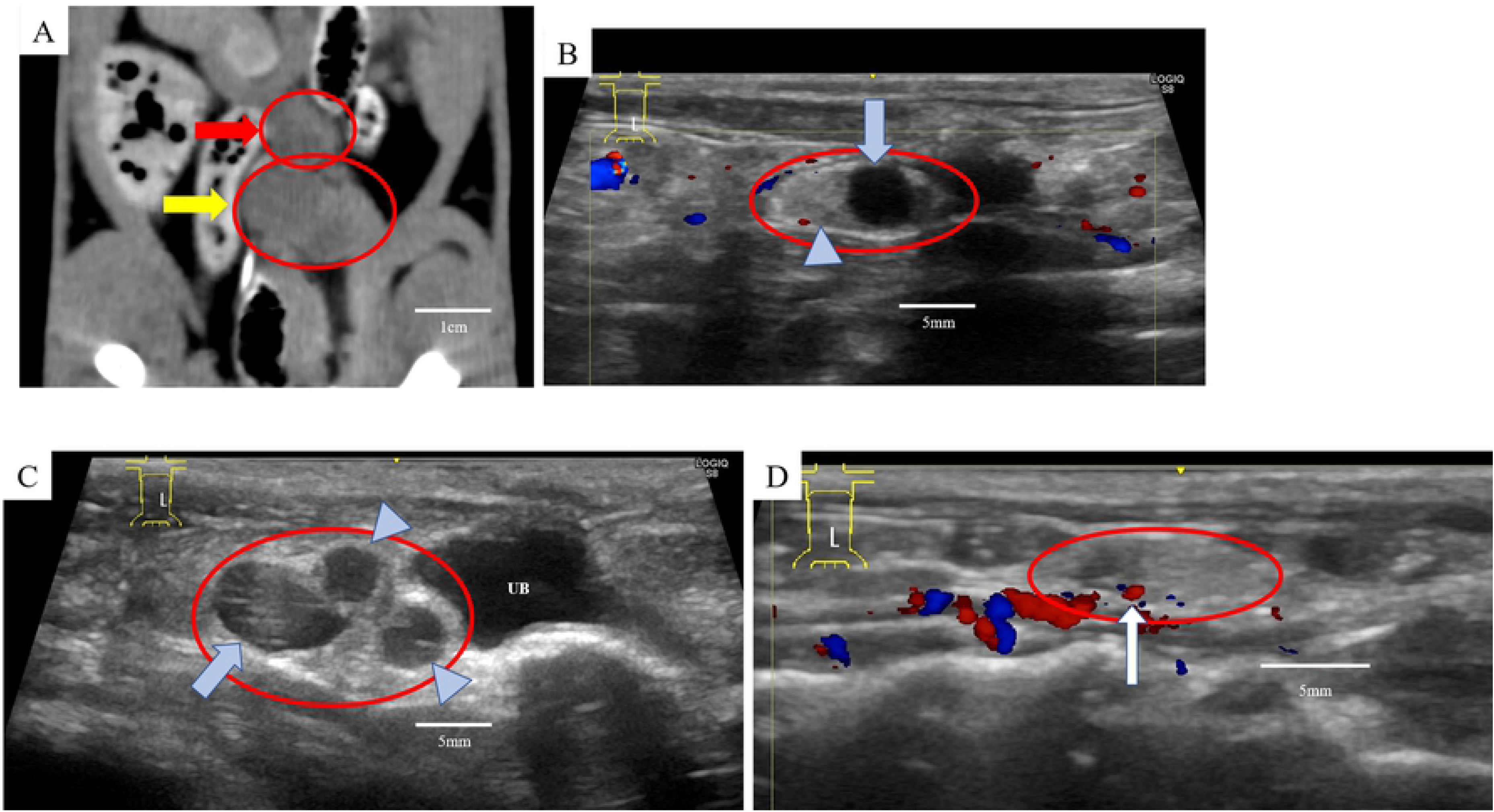
Computed tomography (CT) and ultrasonography images in Experiment 1. A) Visualization of MNB on the abdominal arteriovenous vein under the retroperitoneal area by contrast-enhanced CT. B) Ultrasonographic images of MNB considered to be maturing normally as urine retention was observed in the MNB bladder. C) Ultrasonography image of suspected hydronephrosis of the MNB. D) The blood flow around the MNB could be confirmed by color Doppler using an ultrasound. ※red arrow: MNB1, yellow arrow: MNB2, red circle: MNB, blue arrow: bladder of MNB, arrow head: metanephros of MNB, UB: Recipient’s bladder

**Fig. 2.**
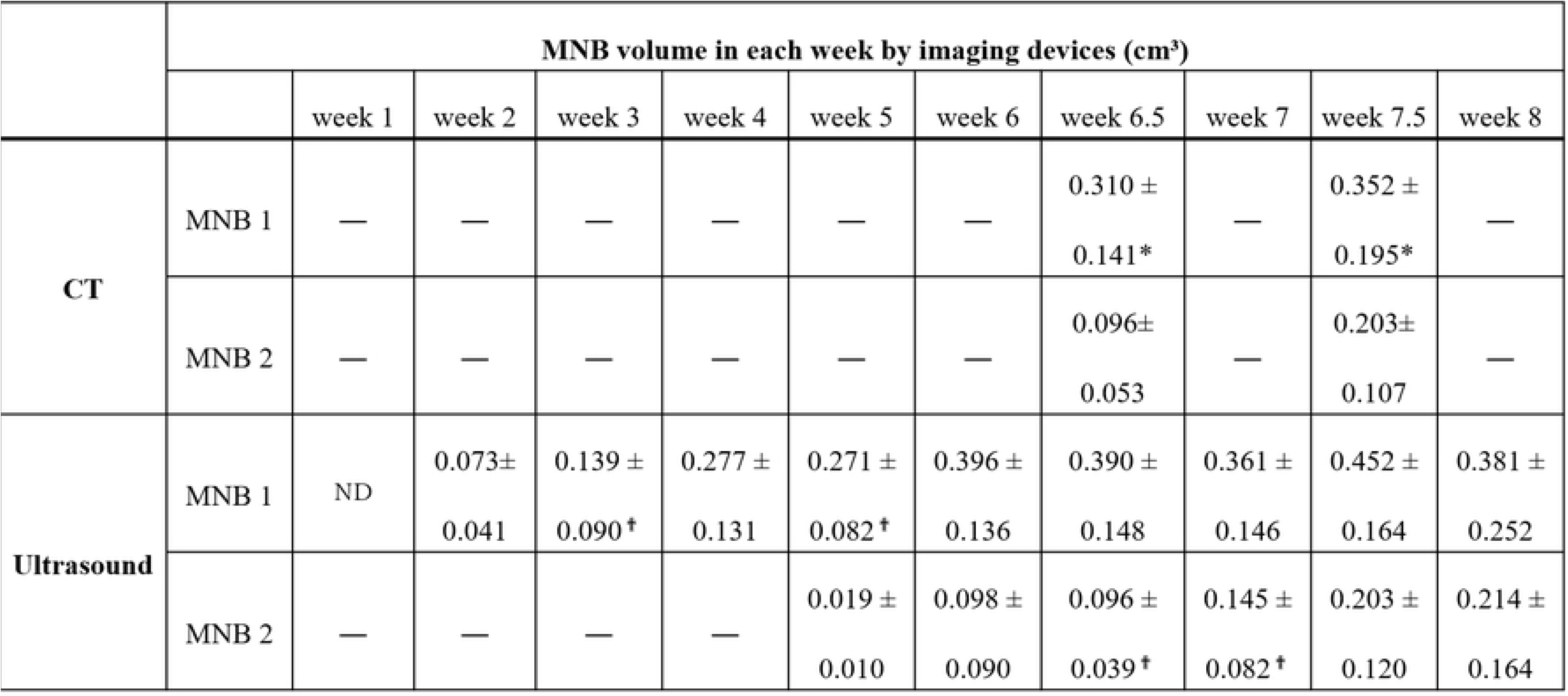
MNB volume transition during observation by contrast-enhanced CT and ultrasonography examinations in Experiment 1. The figure shows the volumes of MNB by CT and ultrasound. The volume of CT was measured at 6.5 and 7.5 weeks and that of ultrasonography was performed weekly until 8 weeks after transplantation of MNB1. \**p* <0.01 vs. ultrasound, † *p* <0.01 vs. the week after. MNB: metanephros with bladder, CT: computed tomography, Week: weeks after transplantation of MNB1.

#### Correlation of volume assessment with urine volume retained in MNB1 and MNB bladder volume assessment

The comparison of volumes measured by contrast CT and ultrasonography examinations (Week 2.5: MNB1, R = 0.78; MNB2, R = 0.79 and Week 3.5: MNB1, R = 0.90; MNB2 = 0.94) indicated a strong positive correlation between MNB1 and MNB2 (Fig. 3). Furthermore, the amount of urine collected from the MNB1 showed a strong positive correlation with the MNB volume determined by ultrasonography (R = 0.89).

**Fig. 3.**
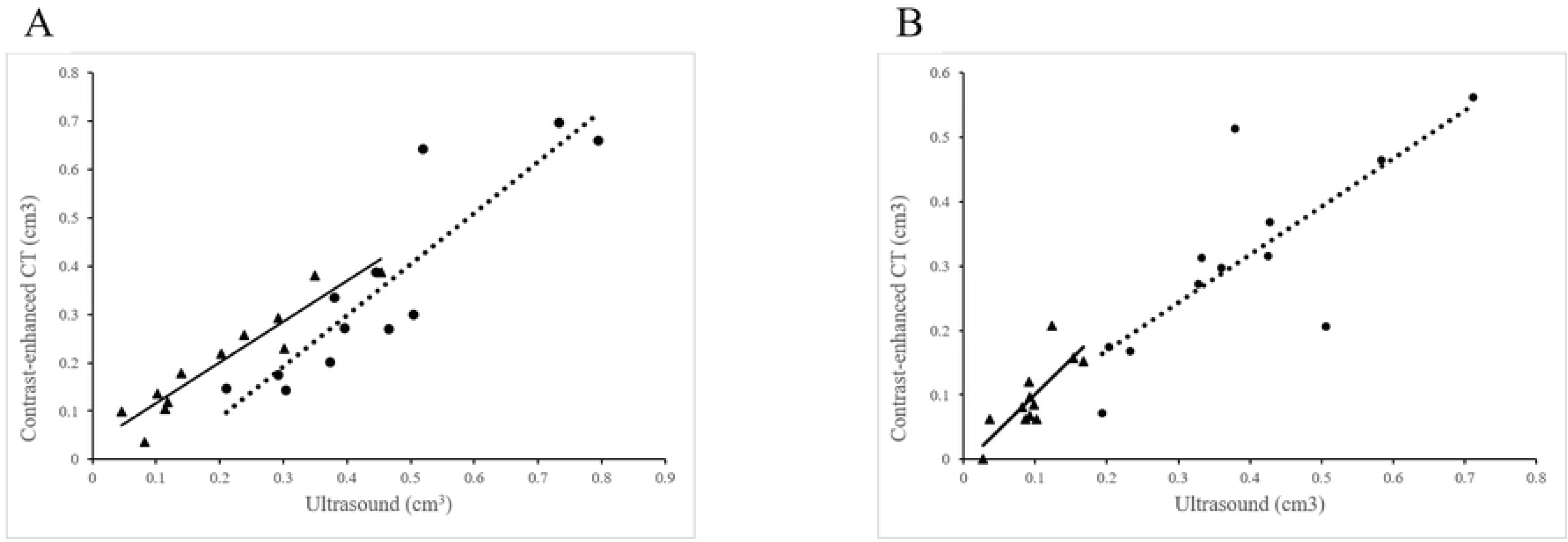
Correlation of the volume between ultrasonography and contrast computed tomography inspection at each week in Experiment 1. The dotted line is the MNB1 and the solid line is the MNB2. A) 6.5 weeks after MNB transplantation (MNB1, R = 0.78; MNB2, R = 0.79). B) 7.5 weeks after MNB transplantation (MNB1, R = 0.90; MNB2, R = 0.94).

#### Tubular dilation and fibrosis in MNB1 and MNB2 by histopathological examinations

Tubular dilation was significantly larger in the non-anastomotic than in the anastomotic group (*p* <0.05). For MNB2, no significant difference was observed regardless of the anastomosis or non-anastomosis of MNB1. However, tubular dilatation was observed in all MNB2. Additionally, there was no significant difference in fibrosis between MNB1 and MNB2, irrespective of anastomosis or non-anastomosis. This indicates that MNB2 shows the same level of growth and induces fibrosis regardless of the degree of renal impairment due to tubular dilation of MNB1.

### Experiment 2

#### Number of days from MNB transplantation to MNB removal

Results are shown in Fig 4. All MNBs could be confirmed by Week 2.5. MNBs with no bladder formation or with hydronephrosis were deemed poorly developed (3/18). The number of days for MNB removal, aside the poorly developed ones, were 20.7 ± 3.6 days (17 to 29 days). For the 72.2% (13/15) of the excised MNB, it was deemed appropriate for urinary tract reconstruction to be performed within 3 weeks after transplantation. For the 20% (3/15), 26.7% (4/15), 40% (6/15), and 13.3% (2/15), 17 days, 19 days, 21 days, and 28 days or more were deemed appropriate timing for reconstruction, respectively.

**Fig. 4.**
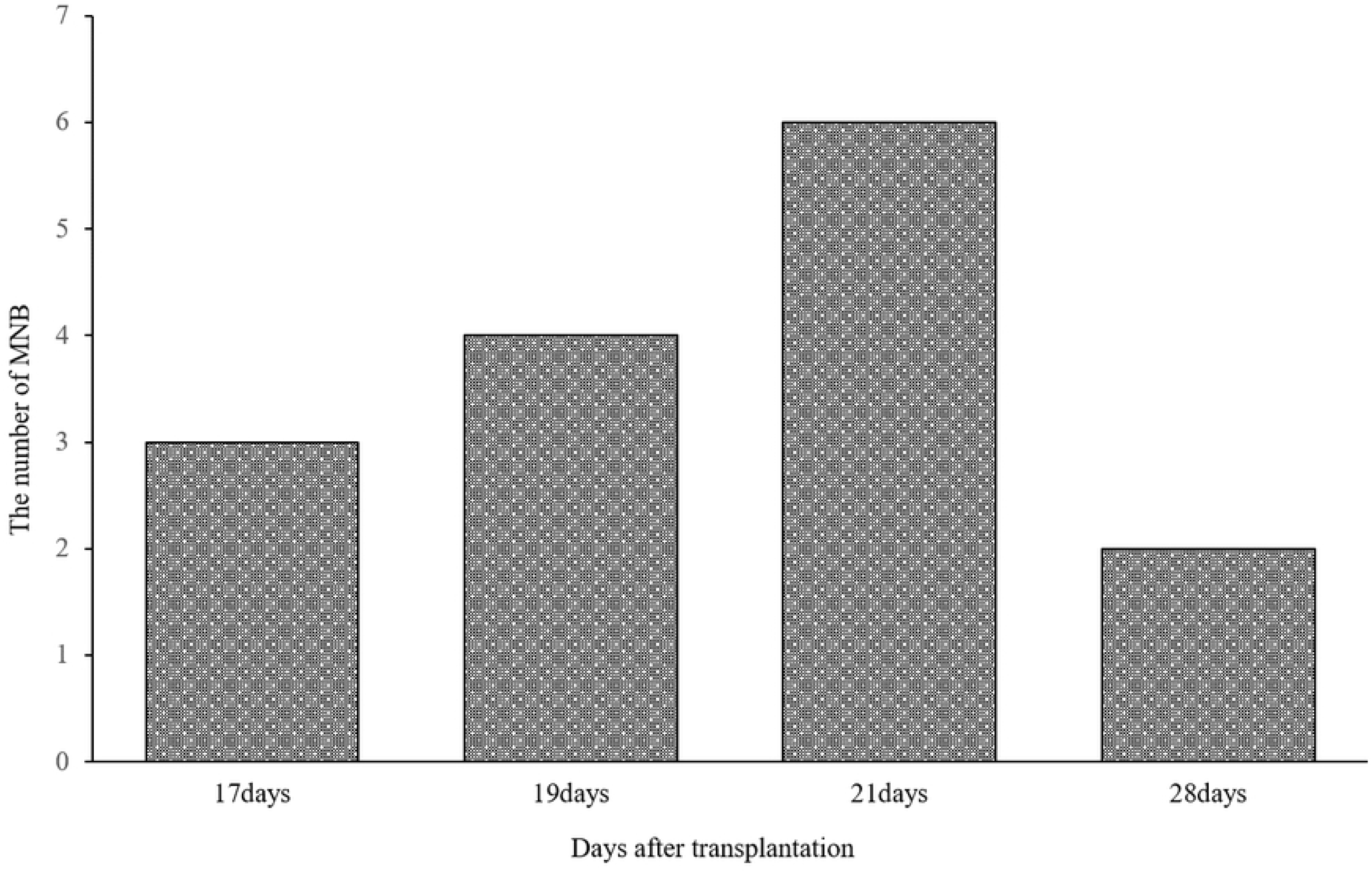
The number of days until the urinary tract reconstruction age and number of MNBs excised after transplantation in Experiment 2. The table shows the number of MNBs excised at a time considered to be an appropriate time for anastomosis. Most of MNBs were removed within 3 weeks after transplantation.

#### Number of days to MNB removal and progression in rate of tubular dilation and fibrosis

The MNBs removed 21 or more days after transplantation had significantly milder tubular dilation than those removed less than 21 days after transplantation (*p* <0.01) (Fig. 5 A). There was no difference in fibrosis in MNB removed 21 days prior and 21 days after transplantation. (Fig. 5 B).

**Fig. 5.**
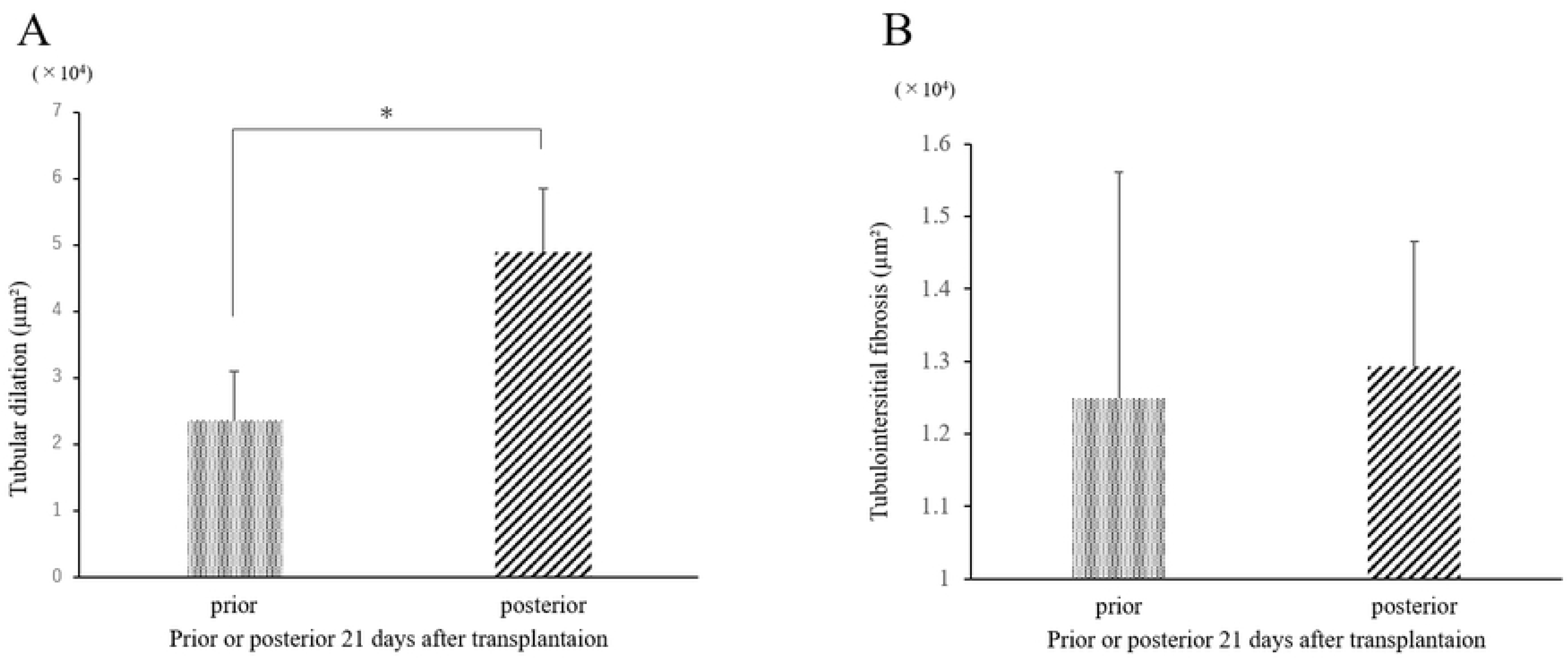
Comparison of tubular dilation and interstitial fibrosis in MNB removed 21 days prior and 21 days after transplantation. A) Tubular dilation, B) Tubulointerstitial fibrosis. The degree of tubular dilation 21 days prior to transplantation decreased significantly compared with that of 21 days after transplantation. * *p* < 0.01

### Experiment 3

#### GFR value of MNB 8 weeks after transplantation

Table 2 shows the GFR measurement results. GFR could be measured in all the animals in the SP and 28UR groups. Although no significant difference was observed between the 28UR and the SP groups, none of the animals in the SP group had a GFR of 0%.

**Table 1.** MNB detection rate by ultrasonography from weeks 1–4 after transplantation in Experiment 1.

|  | Detection rate of MNB |  |  |  |
| --- | --- | --- | --- | --- |
|  | Week 1 | Week 2 | Week 3 | Week 4 |
| MNB 1 | 16.7% | 83.3% | 100% | 100% |
| MNB 2 | 75% | 91.7% | 100% | 100% |

MNB1: the first transplanted MNBs in week 0, MNB2: the second transplanted MNBs in week 4. The detection rates during observation by ultrasound after transplantation in each of MNB1 and MNB2 are shown in the table.

**Table 2.** Comparison of GFR values and normal values of MNB

|  | SP group | 28UR group |
| --- | --- | --- |
| **GFR (ml/min/m<sup>2</sup>)** | 1.95 ± 1.04 | 1.23 ± 1.22 |
| **Compared to Normal**<br>**rats (%)** | 1.1 ~ 5.7% | 0 ~ 5.1% |

#### Tubular dilatation and interstitial fibrosis in MNB of experiment 3 in histopathological examinations

An image of the extracted MNB is shown as an example (Fig. 6 A). In the SP group, the color of the surface of the metanephros could be visually confirmed to have blood flow, and the SP group grew without hydronephrosis. As shown in Fig. 6 B, in the 28UR group, the observed shape of metanephros was irregular. In some cases, the metanephros was hydronephrotic without liquid storage in the MNB bladder. Fig. 7 shows a micrograph of HE staining for the evaluation of tubular dilatation. The 28UR group tended to expand compared to the SP group, but no significant difference was observed between the two groups (Fig. 7 C).

**Fig. 6.**
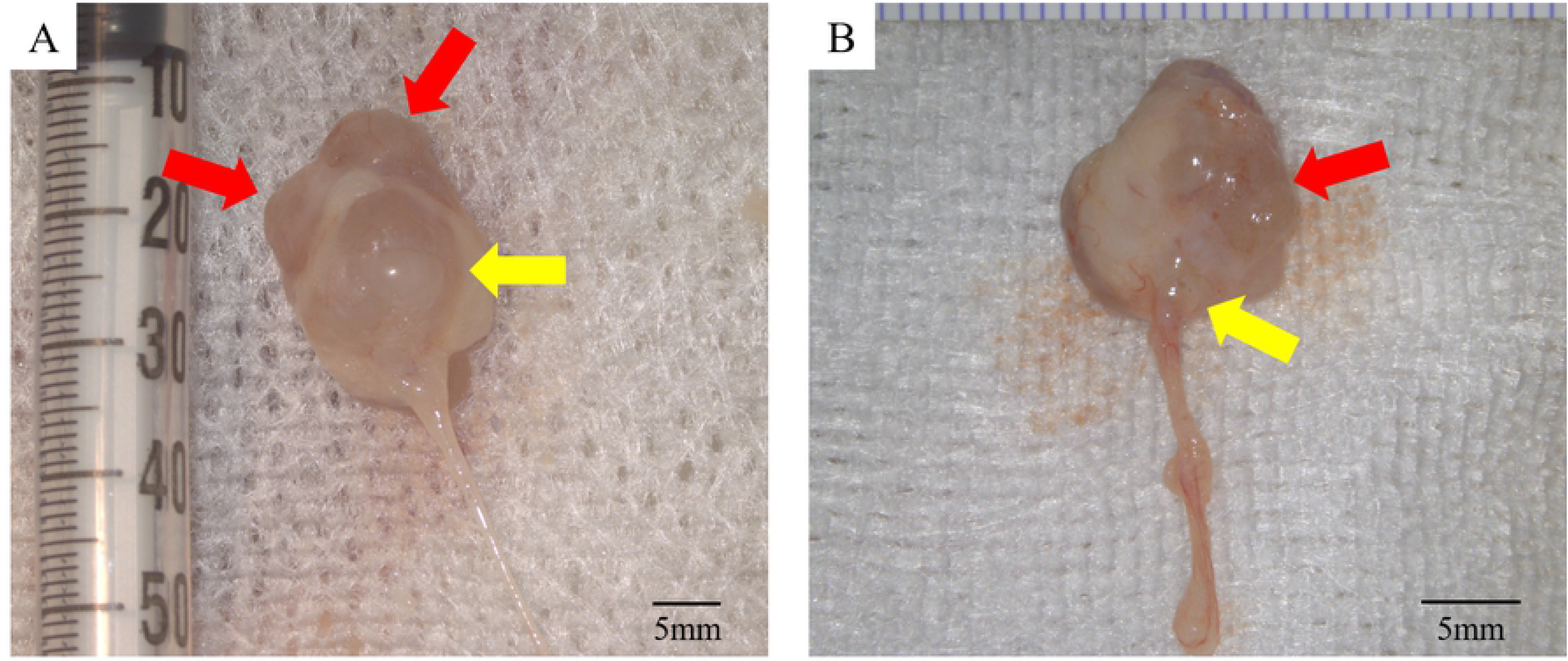
Example of extracted MNB in Experiment 3. The red arrows indicate the metanephros and the yellow arrows indicate the bladder. Fig. 6-A shows that the metanephros did not expand and the MNB is considered to have grown steadily. By contrast, the MNB in the Fig. 6-B has an irregular shape with severe metanephros hydronephrosis.

**Fig 7.**
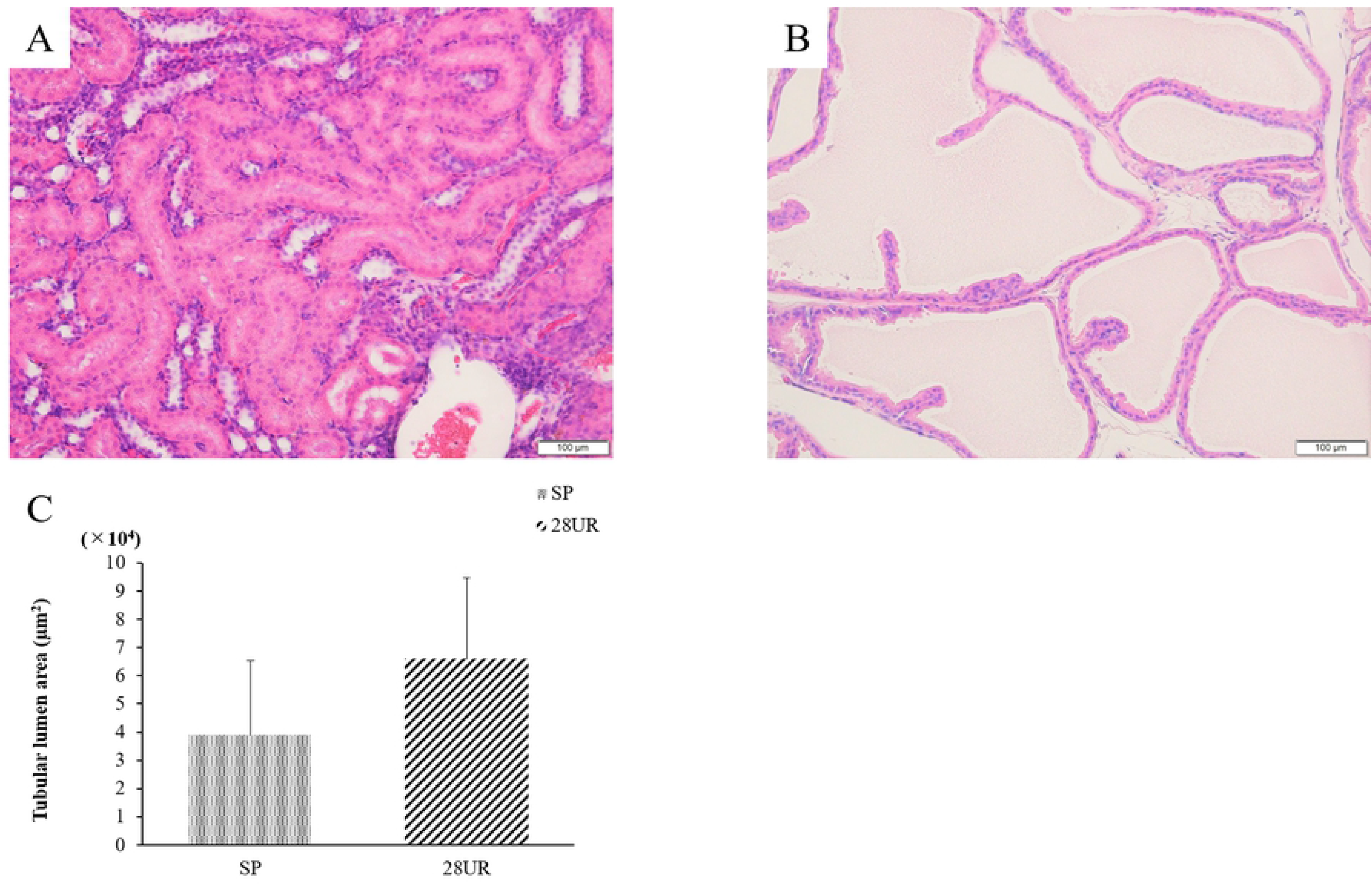
Histopathological examination images of tubular in Experiment 3. A) MNB which was judged to be a suitable period of SWPU method by ultrasonography. B) Histopathology of metanephros with hydronephrosis confirmed by ultrasonography at 4 weeks. C) Comparison of tubular expansion area between the SP and 28UR groups.

MT staining for the evaluation of interstitial fibrosis is shown in Figs. 8 A and B. A comparison of interstitial fibrosis is shown in Fig. 8 C. Interstitial fibrosis was significantly less in the SP than in the 28UR group (*p* <0.01). These results indicated that fibrosis was progressing from the initial stage of tubular dilation.

**Fig. 8.**
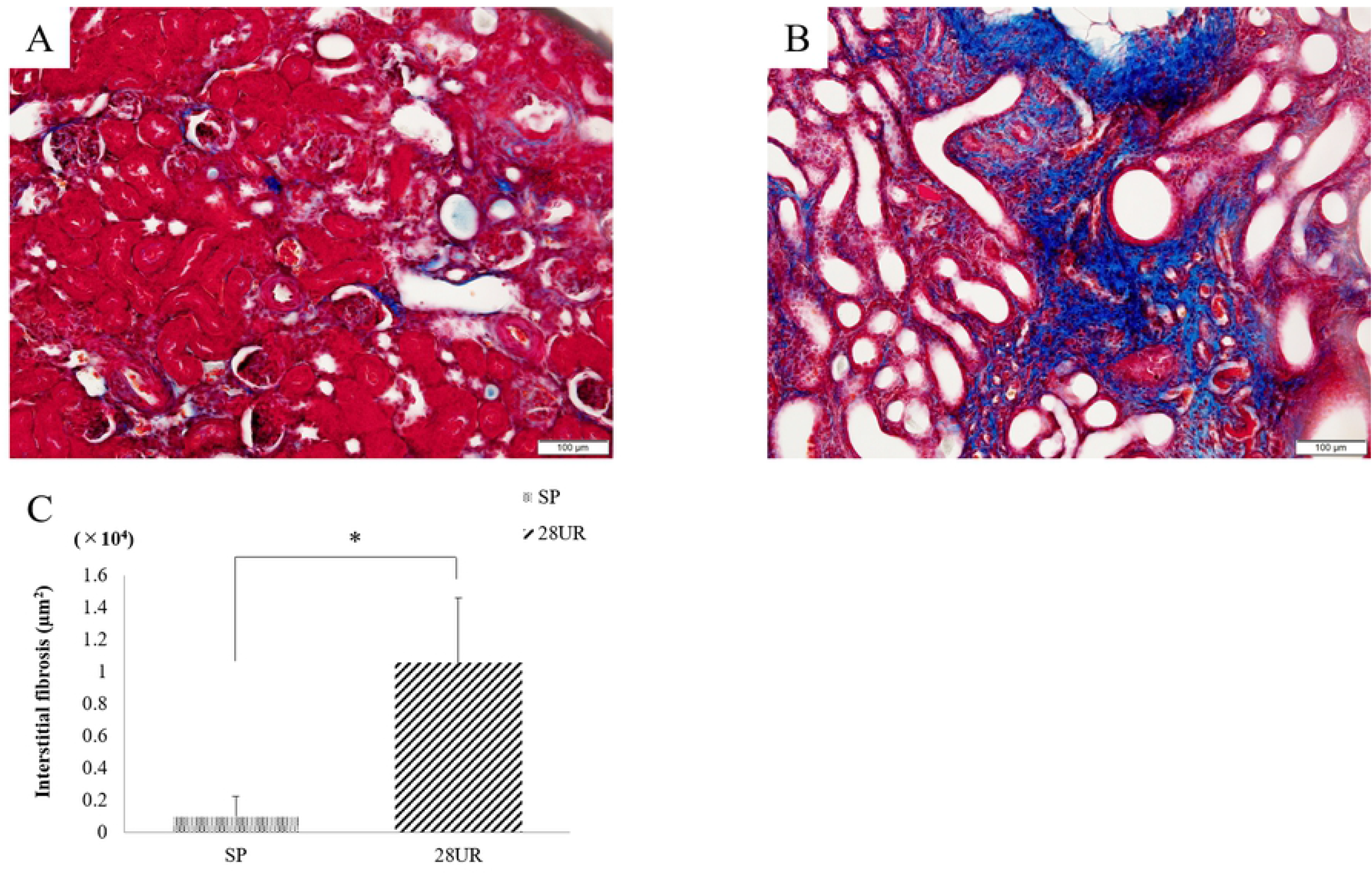
Interstitial fibrosis in MNB in Experiment 3. A) MNB which was judged to be a suitable period of SWPU method by ultrasonography. B) Histopathology of the metanephros with hydronephrosis confirmed by ultrasonography at week 4. C) Comparison of interstitial fibrosis area between the SP and 28UR groups.

The measurements of fibrosis marker are shown in Fig. 9. TGF-β1 was strongly expressed in the 28UR group; mainly in the tubular cells, interstitial, and glomeruli, and similarly, vimentin and type I collagen-α1 showed significant expression in the tubular interstitial. The SP group was significantly milder in all evaluations than the 28UR group (*p* <0.01) (Fig. 9 G).

**Fig. 9.**
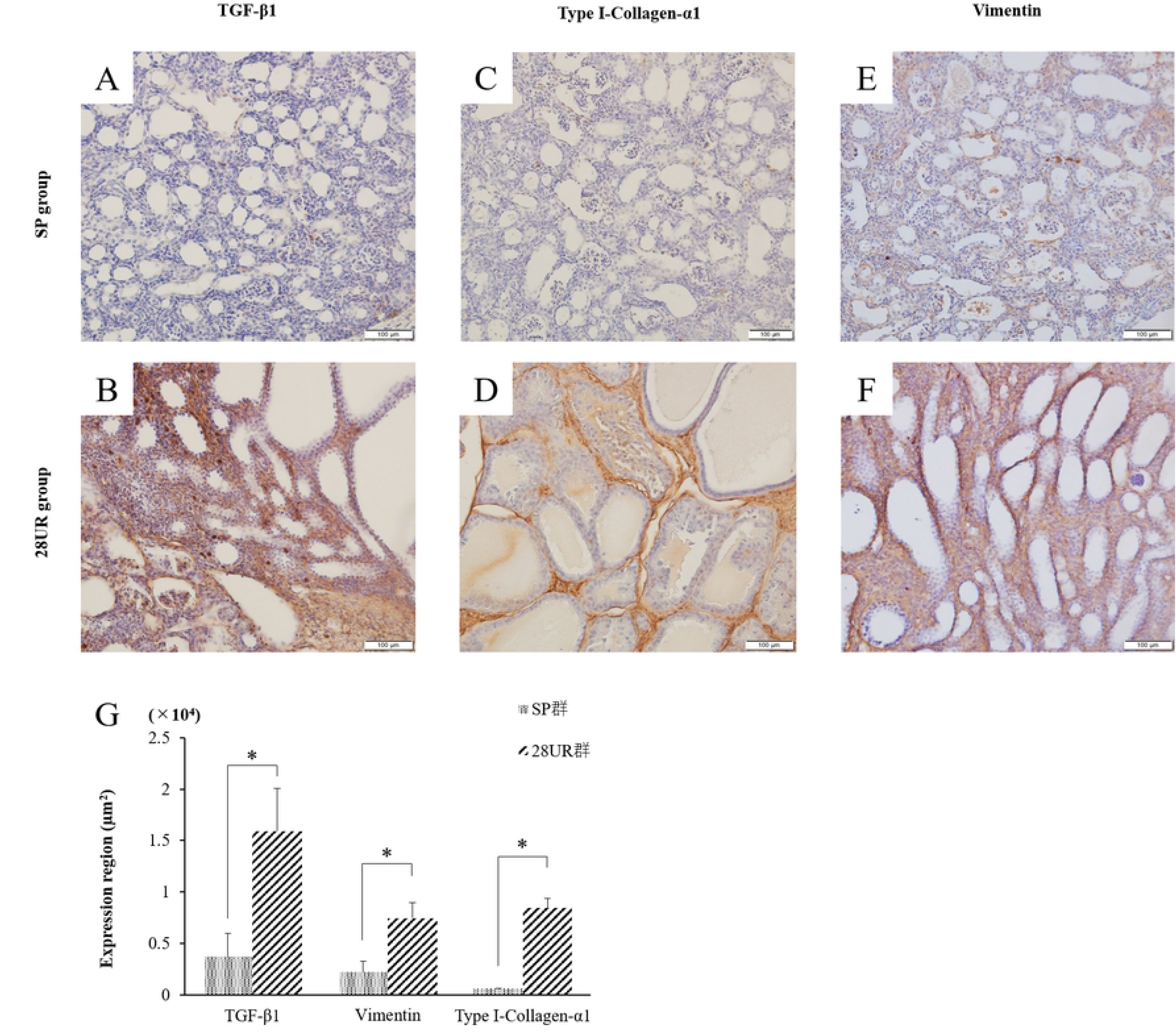
Expression of TGF-β, collagen and vimentin in Experiment 3. A, D) stained image of TGF-*β* 1. B, E) stained image of Type I collagen-*α*1. C, F) stained image of Vimentin. G) All of these assays in the SP group were significantly lower than those of the 28UR group (*p* <0.01).

The image of TUNEL staining and the comparison of the ratio of apoptotic cells in the tubular and glomerular cells are shown in Fig. 10 A. The percentage of apoptotic cells was significantly lower in the SP group than in the 28UR group (*p* <0.05) (Fig. 10 B). Furthermore, there was a strong positive correlation between TGF-β1 expression and the percentage of apoptotic cells in both the glomerular and tubular cells (*p* <0.01) (Figs. 10 C and D).

**Fig. 10.**
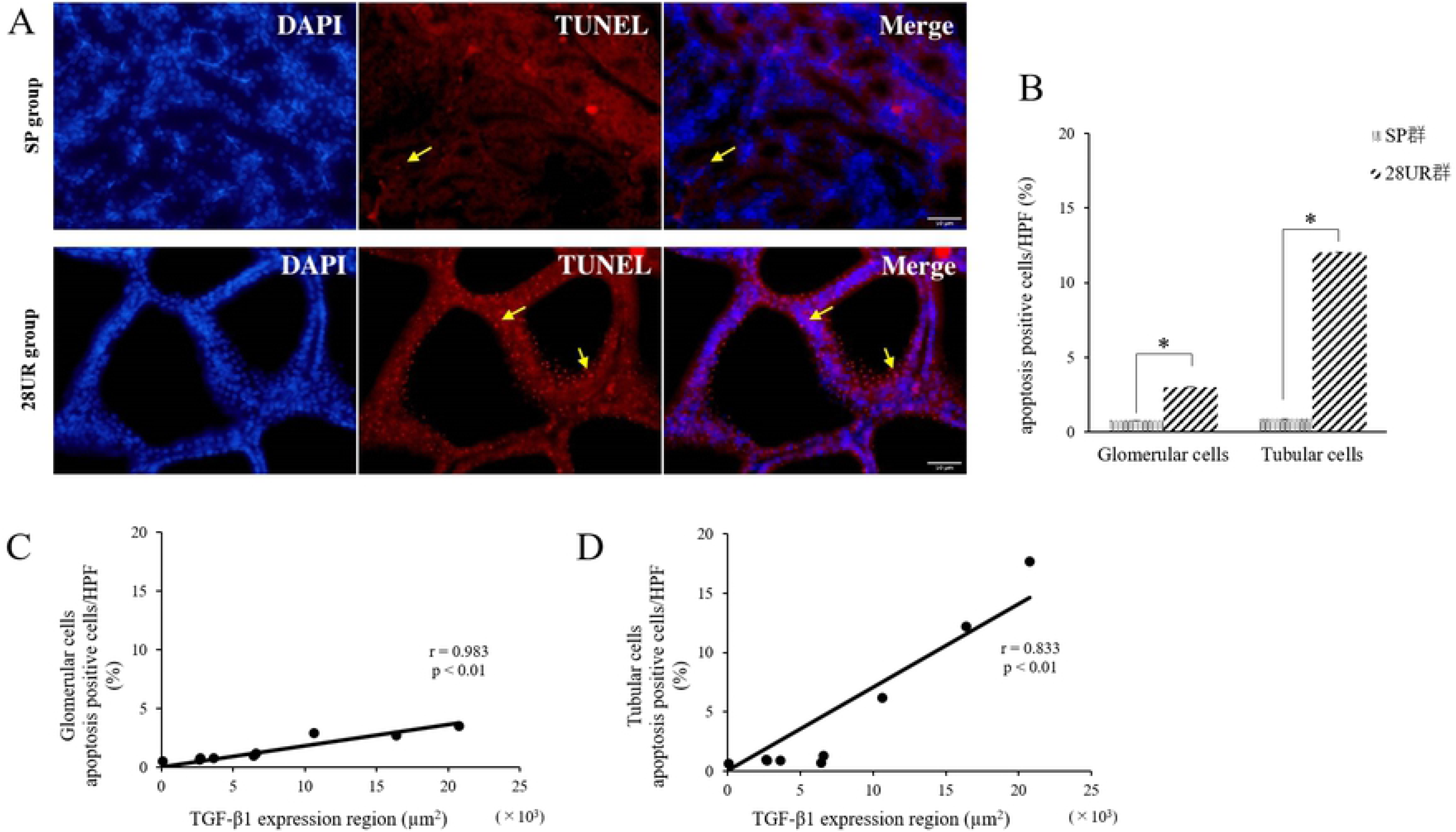
Detection of apoptosis by fluorescent staining, and correlation between expression of TGFβ-1 and apoptosis positive rate. A) Image of TUNEL staining in SP group; Yellow arrows indicate apoptotic cells. B) Image of TUNEL staining in 28UR group; this confirmed a number of apoptotic cells in tubular cells. C) Correlation between apoptosis rate and TGF-β1 expression region in glomerular cells, D) Correlation between apoptosis rate and TGF-β1 expression region in tubular cells

## Discussion

Currently, urinary tract reconstruction or extraction is performed in the transplanted MNB and metanephros approximately 3 to 6 weeks after transplantation, depending on the animal species and the transplantation site [11,19–21]. Metanephros weight gain stops at about 4 weeks after the transplantation [2], and it has been reported that after development, the GFR is about 3%–11% of that of normal kidneys, as the metanephros alone is a small tissue [21,22]. However, the survival time did not differ from the cases in which one MNB was transplanted, even in experiments in which several MNBs were transplanted [13]. One of the factors is that the metanephros has a remarkable degree of hydronephrosis and fibrosis. Obstruction during the development of rat kidneys has been reported to cause developmental suppression and persistent damage after maturation [14,15]. For this reason, it was necessary to investigate the appropriate timing for SWPU. Therefore, as in previous papers, two MNBs were transplanted, and their development was closely observed by ultrasonography and CT.

Contrast-enhanced CT showed no enhancement of the metanephros and bladder of MNB; however, it was possible to evaluate the MNB to the extent that the MNB volume was measured, and this was regarded as an accurate volume index. A single intravenous dose of iohexol, the nonionic iodine-based contrast agent, undergoes rapid clearance from the blood of rats and translocates to the tissues [23]. It rapidly migrates to the kidneys and is distributed at high concentrations [23]. Yokote et al. [11] previously performed contrast-enhanced CT on MNB to confirm urinary patency after SWPU in MNB. In that study, both kidneys were removed, and the anastomotic ureter was ligated; angiography was performed to visualize the recipient ureter [11]. In this present study, we considered that one of the recipient’s kidneys was present, resulting in the excretion of the contrast agent before it flowed into the MNB.

Ultrasonography detected MNB in all animals, similar to contrast-enhanced CT examinations. Ultrasonography assessed MNB morphologically, unlike contrast-enhanced CT. As the MNB was transplanted into the retroperitoneum, it was possible to be identified at an earlier stage than expected by specifying the expansion of the retroperitoneal cavity. We also observed blood flow to the MNB using ultrasonography and observed the neovascularized vessels. Urine production has been previously confirmed by metanephros transplantation or MNB transplantation [11,12,24]. Therefore, as we hypothesized, the MNB bladder was visualized using low echo, making retrieval easier. In children with congenital hydronephrosis, ultrasonography can reveal septum and cyst in the kidneys when severe hydronephrosis occurs [25,26]. Here, it was also possible to evaluate hydronephrotic metanephros without a urinary excretion pathway, because the parenchyma was visualized as a vacuole with an indistinct parenchyma, like severe hydronephrosis in a developed kidney. Spheroid volume measurements by ultrasonography was also used in the kidney, thyroid, and prostate [27,28–31]. As the MNB volume determined by ultrasonography in this study strongly correlated with that of the CT, and the MNB was approximated despite the very small volume, volume calculation using the spheroid equation was also considered very accurate for MNB. Therefore, ultrasonography is considered to be a simple, minimally invasive, and useful method, considering the effects on the contrast medium and radiation as in contrast-enhanced CT examinations.

In Experiment 1, there was no difference in tubule dilation and fibrosis in MNB2 compared with MNB1, regardless of the presence or absence of SWPU in MNB1. This suggests that, by 4 weeks after transplantation, many metanephros had already experienced hydronephrosis, and the appropriate timing had passed with or without urinary tract reconstruction. Indeed, previous reports referred to the presence of MNBs that had experienced hydronephrosis by 3 weeks after transplantation [11]. A comparison of the growth rates between MNB1 and MNB2 showed no significant difference in the MNB volumes, but rather showed that the detection rate by ultrasonography was higher in the first week of MNB2. A plurality of blood vessels around the MNB1 and MNB2 were confirmed using a color Doppler method for the first 2 weeks after transplantation. It has been confirmed that the transplanted metanephros regenerates recipient-derived blood vessels and is chimerized with the donor’s metanephros [32]. Angiogenesis involves factors such as vascular endothelial growth factor, platelet-derived growth factor, and fibroblast growth factor [33]. These angiogenic factors play important roles in tissue ischemia and angiogenesis. One report suggests that the metanephros is provided with a spatial direction for capillary development by VEGF [34] and may be involved in the vasculature identified around the MNB. However, it must be considered that VEGF is also involved in fibrosis, which promotes the growth of grafts and may further exacerbate fibrosis during hydronephrosis [35,36]. In the unilateral ureteric obstruction (UUO) model of progressive injury, an angiogenic response was observed early and the endothelial cells proliferated; however, this led to endothelial cell loss after the 4^th^ day [37]. Therefore, the neovascularized capillaries disappeared, and the tissue fell into an ischemic state again. The high detection rate of the MNB2 and the presence of tubular dilation at 4 weeks after transplantation suggest that the neovascularization of the MNB1 may also affect MNB2. The MNB2 could be affected by MNB1 angiogenesis because it was implanted just above the MNB1 in the retroperitoneal cavity. One of the reasons may be that one kidney was removed during the MNB2 transplantation. Studies in fetal ewes have shown that in the event of a sudden loss of unilateral renal function due to obstruction in the unilateral ureter, the remaining kidney plays a compensatory role, resulting in kidney enlargement [38]. Moreover, a growth period of 4 weeks may be sufficient to develop into hydronephrotic metanephros of MNB2. This suggests that the degree of growth was not constant when multiple grafts were transplanted and may vary greatly between MNBs.

In Experiment 2, only one MNB was transplanted, and the MNB growth rate and appearance of MNB bladder were observed over time in more details to establish an index of the appropriate timing of SWPU. Even when only one MNB was transplanted, the growth rate varied among individuals as can be inferred from the results. Particularly well-developed MNB showed bladder dilation within the 3 weeks after transplantation with a percentage of 38.9% of the total. By week 3 after transplantation, more than half of the MNB had the appropriate timing for urinary tract reconstruction, suggesting that SWPU should be done earlier than previously reported. Furthermore, there was a poorly developed MNB without appearance or change in size of the MNB, bladder from the third week onward. Histopathological examination did not show excessive tubular dilation and progression of fibrosis as observed in Experiment 1 in MNB resected at the appropriate stage of urinary tract reconstruction. Obstruction causes increased ureteral/pelvic pressure and tubular dilatation in the kidney. This increase in pressure stimulates tubular epithelial cells and causes fibrosis to progress due to epithelial-mesenchymal transition [39]. Previous studies, using the rat UUO model, showed the expression of TGF-β1 immediately after the obstruction of the urinary tract and the expression of vimentin and myofibroblasts 2 days after the obstruction [37]. The same is true for the experiment conducted in the organogenesis stage using the rat neonatal UUO model, and the growth rate decreased by 30% even after unblocking 5 days after obstruction, while all of blood pressure, GFR, urine flow, and sodium/potassium excretion decreased [15]. Therefore, it has been shown that UUO is particularly susceptible to long-term effects immediately after kidney formation [14,15]. Although the transplanted MNB cannot be completely explained by the neonatal UUO model, considering the period after transplantation as being in the neonatal period, it can be expected that the occlusion period is similarly involved in the growth of the transplanted metanephros. Here, ultrasonographic observations were performed every other day; some animals showed rapid bladder dilation even on this day. Rats have only 10% of the nephron formed at birth, and it is said that kidney formation is completed within the first week after birth [15]. The fetus used this time was 15 days of fetal age, and the birth of the rat is usually 21 to 23 days of fetal age on average; therefore, considering that the morphological formation of the kidney is completed 2 weeks after transplantation, a rapid development of urine in the MNB bladder, 2 to 3 weeks after transplantation is considered as normal development. It can be said that, even in the case of MNB, those that grew well may grow at the same growth rate as the organs in the fetal rat body that grew normally. For this reason, it is important to observe and evaluate MNB bladder dilation daily from the second week when MNB grows rapidly. GFR measurement had been performed, when the metanephros was transplanted, and showed a low value of 3% to 11% of normal. Although not the same measurement method, the value of 5% of normal in the SP group in this study was within the range we assumed. This is because, in previous reports, the metanephros grew about 90 to 116 days after reconstructing the urinary tract [24], and it is thought that a certain amount of tissue was present. Urinary tract reconstruction for these metanephros was performed mainly at a time determined by naked eye and may have been exposed to long-term obstruction. Prolonged obstruction results in the persistent dilation of the tubule, the glomerular Bowman’s capsule, and the production of inflammatory cytokines such as TGF-β from the tubular cells [40]. TGF-β1 is expressed in tissues such as the hematopoietic tissue, endothelial tissue, and bone tissue in developing embryos, and acts as an important growth factor as in heart formation [41,42]. However, TGF-β1 is also deeply involved in fibrosis, causing epithelial to mesenchymal transition (EMT) in tubular cells in the kidney and inducing expression of vimentin, collagen, and alpha-smooth muscle actin [40]. In addition, TGF-β1 targets protease inhibitor-1 (PAI-1) in proximal tubular epithelial cells and stromal fibroblasts, and the transcription factor p53 replicates through this PAI-1. It causes an aging state and induces cell growth inhibition and apoptosis [40,43–45]. TGF-β1 expression has also been reported in the metanephros in fetuses and our defined urinary tract reconstruction at the appropriate anastomosis stage in MNB inhibited TGF-β1 expression due to persistent inhibition of tubular dilatation. It is presumed that the replacement suppressed tissue replacement by collagen and vimentin. Therefore, the fact that GFR could be measured in tissues, approximately, 60 days after transplantation would be a great knowledge.

There are some limitations to this study. First, we only performed diagnostic imaging-based assessments and did not assess aspects such as renin and erythropoietin activities. Second, because we assessed the MNB only for a short period (60 days after transplantation), we did not perform long-term assessment of function and morphology of the transplants. These issues need to be elucidated further in future studies.

In conclusion, this is the first study to successfully observe the time course of MNB in detail by ultrasonography. The appropriate timing for urinary tract reconstruction in rats by SWPU, as revealed by ultrasonography, was when the metanephros has not undergone hydronephrosis, and the early period from when there was urinary retention of 0.016 cm^3^ in the MNB bladder. This method suppressed the excessive dilatation of the renal tubules of the transplanted metanephros, thereby providing evidence to reduce the progression of fibrosis. We believe that this will greatly contribute to the evaluation of MNB development and urinary tract reconstruction in xenotransplantation and in human clinical practice. In addition, there is a possibility that it can be found even when transplanted to another site. It is also important to evaluate the morphology and function of small grafts in the retroperitoneal cavity while minimizing invasion when considering transplantation into patients in the future.

## Acknowledgments

No third-party funding or support was received in connection with this study or the writing or publication of the manuscript. We would like to thank Dr. Satoshi Kameshima for advice to experiment and Editage (www.editage.com) for English language editing.

## Supporting information

**S1 Fig. Stepwise peristaltic ureter (SWPU) system**

The recipient is transplanted with the MNB and allowed to grow. The ureter of the recipient is anastomosed to the MNB bladder where urine has accumulated. In this manner, MNB can measure urinary excretion.

**S2 Fig. Experimental procedure**

A) Experiment 1, B) Experiment 2, C) Experiment 3

**S Table. GFR measurements in healthy adult rats using inulin clearance**

Measurements are performed in the same situation as the GFR measurement method in the MNB.

## References

1. Hill NR, Fatoba ST, Oke JL, Hirst JA, O’Callaghan CA, Lasserson DS, et al. Global Prevalence of Chronic Kidney Disease - A Systematic Review and Meta-Analysis. PLoS One. 2016;11:e0158765.1

2. Mills KT, Xu Y, Zhang W, Bundy JD, Chen CS, Kelly TN, et al. A systematic analysis of worldwide population-based data on the global burden of chronic kidney disease in 2010. Kidney Int. 2015;88:950–957.

3. Takasato M, Er PX, Chiu HS, Maier B, Baillie GJ, Ferguson C, et al. Kidney Organoids From Human iPS Cells Contain Multiple Lineages and Model Human Nephrogenesis. Nature. 2015;526:564–568

4. Taguchi A, Kaku Y, Ohmori T, Sharmin S, Ogawa M, Sasaki H, et al. Redefining the in Vivo Origin of Metanephric Nephron Progenitors Enables Generation of Complex Kidney Structures from Pluripotent Stem Cells. Cell Stem Cell. 2014;14,53–67.

5. Yokoo T, Ohashi T, Shen JS, Sakurai K, Miyazaki Y, Utsunomiya Y, et al. Human mesenchymal stem cells in rodent whole-embryo culture are reprogrammed to contribute to kidney tissues. Proc Natl Acad Sci U S A. 2005;102:3296–3300.

6. Matsumoto K, Yokoo T, Matsunari H, Iwai S, Yokote S, Teratani T, et al. Xenotransplanted embryonic kidney provides a niche for endogenous mesenchymal stem cell differentiation into erythropoietin-producing tissue. Stem Cells. 2012;30:1228–1235.

7. Yokoo T, Fukui A, Ohashi T, Miyazaki Y, Utsunomiya Y, Kawamura T, et al. Xenobiotic Kidney Organogenesis from Human Mesenchymal Stem Cells Using a Growing Rodent Embryo. J Am Soc Nephrol. 2006;17:1026–1034.

8. Yamanaka S, Tajiri S, Fujimoto T, Matsumoto K, Fukunaga S, Kim BS, et al. Generation of interspecies limited chimeric nephrons using a conditional nephron progenitor cell replacement system. Nat Commun. 2017;8:1719.

9. Fujimoto T, Yamanaka S, Tajiri S, Takamura T, Saito Y, Matsumoto K, et al. In vivo regeneration of interspecies chimeric kidneys using a nephron progenitor cell replacement system. Sci Rep. 2019;9:6965.

10. Tajiri S, Yamanaka S, Fujimoto T, Matsumoto K, Taguchi A, Nishinakamura R, et al. Regenerative potential of induced pluripotent stem cells derived from patients undergoing haemodialysis in kidney regeneration. Sci Rep. 2018;8:14919.

11. Yokote S, Matsunari H, Iwai S, Yamanaka S, Uchikura A, Fujimoto E, et al. Urine excretion strategy for stem cell-generated embryonic kidneys. Proc Natl Acad Sci U S A. 2015;112:12980–12985.

12. Fujimoto E, Yamanaka S, Kurihara S, Tajiri S, Izuhara L, Katsuoka Y, et al. Embryonic kidney function in chronic renal failure model in rodents. Clin Exp Nephrol. 2017;21:579–588.

13. Yokoo T, Fukui A, Matsumoto K, Kawamura T. Kidney regeneration by xeno-embryonic nephrogenesis. Med Mol Morphol. 2008;41:5–13.

14. Chevalier RL, Kim A, Thornhill BA, Wolstenholme JT. Recovery following relief of unilateral ureteral obstruction in the neonatal rat. Kidney Int. 1999;55:793–807.

15. Chevalier RL, Thornhill BA, Chang AY, Cachat F, Lackey A. Recovery from release of ureteral obstruction in the rat: relationship to nephrogenesis. Kidney Int. 2002;61:2033–2043.

16. Truong LD, Gaber L, Eknoyan G. Obstructive uropathy. Contrib Nephrol. 2011;169:311–326.

17. Ichikawa I, Kuwayama F, Pope JC 4th, Stephens FD, Miyazaki Y. Paradigm shift from classic anatomic theories to contemporary cell biological views of CAKUT. Kidney Int. 2002;61:889–898.

18. Kanda Y. Investigation of the freely available easy-to-use software ‘EZR’ for medical statistics. Bone Marrow Transplant. 2013;48:452–458.

19. Marshall D, Clancy M, Bottomley M, Symonds K, Brenchley PE, Bravery CA. Transplantation of metanephroi to sites within the abdominal cavity. Transplant Proc. 2005;37:194–197.

20. Vera-Donoso CD, García-Dominguez X, Jiménez-Trigos E, García-Valero L, Vicente JS, Marco-Jiménez F. Laparoscopic transplantation of metanephroi: A first step to kidney xenotransplantation. Actas Urol Esp. 2015;39:527–534.

21. Rogers SA, Lowell JA, Hammerman NA, Hammerman MR. Transplantation of developing metanephros into adult rats. Kidney Int. 1998;54:27–37.

22. Marshall D, Clancy M, Bottomley M, Symonds K, Brenchley PE, Bravery CA. Transplantation of metanephros to sites within the abdominal cavity. Transplant Proc. 2005;37:194–197.

23. Nagai E, Ishigaki N, Hakusui H. Pharmacokinetic Studies of Iohexol after a Single Intravenous Injection in Rats. Prog Med. 1986;6:2378–2389.

24. Clancy MJ, Marshall D, Dilworth M, Bottomley M, Ashton N, Brenchley P. Immunosuppression is essential for successful allogeneic transplantation of the metanephros. Transplantation. 2009;88:151–159.

25. Isaksen CV, Eik-Nes SH, Blaas HG, Torp SH. Fetuses and infants with congenital urinary system anomalies: correlation between prenatal ultrasound and postmortem findings. Ultrasound Obstet Gynecol. 2000;15:177–185.

26. Fernbach SK, Maizels M, Conway JJ. Ultrasound grading of hydronephrosis: introduction to the system used by the Society for Fetal Urology. Pediatr Radiol. 1993;23:478–480.

27. Hyun CK, Dal MY, Sang HL, Yong DC. Usefulness of Renal Volume Measurements Obtained by a 3-Dimensional Sonographic Transducer with Matrix Electronic Arrays. J Ultrasound Med. 2008;27:1673–1681.

28. Marcello M, Pier P, Antonio S, Raffaele L, Ernesto S, Massimo S, et al. Accuracy of Sonographic Volume Measurements of Kidney Transplant. J Clin Ultrasound. 2006;34:184–189.

29. Noto H, Kizu N, Sugaya K, Nishizawa O, Harada T, Tsuchida T. Prostate measurement by transabdominal ultrasonography. Journal of the Japanese Urology. 1987;78:1071–1076.

30. Reid T, Stacy AL, Stephen RW, Gregory BD. Estimation of feline renal volume using computed tomography and ultrasound. Vet Radiol Ultrasound. 2012;54:127–132.

31. Barberet V, Baeumlin Y, Taeymans O, Duchateau L, Peremans K, van Hoek I, Daminet S, Saunders JH. Pre-and posttreatment ultrasonography of the thyroid gland in hyperthyroid cats. Vet Radiol Ultrasound. 2010;51:324–330.

32. Yokoo T, Fukui A, Matsumoto K, Ohashi T, Sado Y, Suzuki H, et al. Generation of a transplantable erythropoietin-producer derived from human mesenchymal stem cells. Transplantation. 2008;85:1654–1658.

33. Distler JH, Hirth A, Kurowska-Stolarska M, Gay RE, Gay S, Distler O. Angiogenic and angiostatic factors in the molecular control of angiogenesis. Q J Nucl Med. 2003;47:149–161.

34. Tufró A. VEGF spatially directs angiogenesis during metanephric development in vitro. Dev Biol. 2000;227:558–566.

35. Chaudhary NI, Roth GJ, Hilberg F, Müller-Quernheim J, Prasse A, Zissel G, et al. Inhibition of PDGF, VEGF and FGF signalling attenuates fibrosis. Eur Respir J. 2007;29:976–985.

36. Kariya T, Nishimura H, Mizuno M, Suzuki Y, Matsukawa Y, Sakata F, et al. TGF-ß1-VEGF-A pathway induces neoangiogenesis with peritoneal fibrosis in patients undergoing peritoneal dialysis. Am J Physiol Renal Physiol. 2018;314:F167–180.

37. Lin SL, Chang FC, Schrimpf C, Chen YT, Wu CF, Wu VC, et al. Targeting endothelium-pericyte cross talk by inhibiting VEGF receptor signaling attenuates kidney microvascular rarefaction and fibrosis. Am J Pathol. 2011;178:911–923.

38. Peters CA, Gaertner RC, Carr MC, Mandell J. Fetal compensatory renal growth due to unilateral ureteral obstruction. J Urol. 1993;150:597–600.

39. Moeller BJ, Cao Y, Vujaskovic Z, Li CY, Haroon ZA, Dewhirst MW. The relationship between hypoxia and angiogenesis. Semin Radiat Oncol. 2004;14:215–221.

40. Meng XM, Nikolic-Paterson DJ, Lan HY. TGF-β: the master regulator of fibrosis. Nat Rev Nephrol. 2016;12:325–338.

41. Akhurst RJ, Lehnert SA, Faissner A, Duffie E. TGF beta in murine morphogenetic processes: the early embryo and cardiogenesis. Development. 1990;108:645–656.

42. Gatherer D, Ten Dijke P, Baird DT, Akhurst RJ. Expression of TGF-beta isoforms during first trimester human embryogenesis. Development. 1990;110:445–460.

43. Dockrell ME, Phanish MK, Hendry BM. Tgf-beta auto-induction and connective tissue growth factor expression in human renal tubule epithelial cells requires N-ras. Nephron Exp Nephrol. 2009;112:e71–79.

44. Higgins SP, Tang Y, Higgins CE, Mian B, Zhang W, Czekay RP, et al. TGF-ß1/p53 signaling in renal fibrogenesis. Cell Signal. 2018;43:1–10.

45. Schuster N, Krieglstein K. Mechanisms of TGF-beta-mediated apoptosis. Cell Tissue Res. 2002;307:1–14.

